# Applying Super-Resolution and Tomography Concepts to Identify Receptive Field Subunits in the Retina

**DOI:** 10.1101/2023.11.27.568854

**Authors:** Steffen Krüppel, Mohammad H. Khani, Helene M. Schreyer, Shashwat Sridhar, Varsha Ramakrishna, Sören J. Zapp, Matthias Mietsch, Dimokratis Karamanlis, Tim Gollisch

## Abstract

Spatially nonlinear stimulus integration by retinal ganglion cells lies at the heart of various computations performed by the retina. It arises from the nonlinear transmission of signals that ganglion cells receive from bipolar cells, which thereby constitute functional subunits within a ganglion cell’s receptive field. Inferring these subunits from recorded ganglion cell activity promises a new avenue for studying the functional architecture of the retina. This calls for efficient methods, which leave sufficient experimental time to leverage the acquired knowledge. Here, we combine concepts from super-resolution microscopy and computed tomography and introduce super-resolved tomographic reconstruction (STR) as a technique to efficiently stimulate and locate receptive field subunits. Simulations demonstrate that this approach can reliably identify subunits across a wide range of model variations, and application in recordings of primate parasol ganglion cells validates the experimental feasibility. STR can potentially reveal comprehensive subunit layouts within less than an hour of recording time, making it ideal for online analysis and closed-loop investigations of receptive field substructure in retina recordings.

## Introduction

Retinal ganglion cells, the output neurons of the retina, are classically modelled with a linear-nonlinear (LN) model (Chichilnisky, 2001). This can take the center-surround structure of their receptive fields into account, but indiscriminately considers luminance signals inside the receptive field to be integrated linearly and passed through a nonlinearity only afterwards, at the model’s output stage. While the LN model is still popular due to its simplicity, it has long been known that many ganglion cells respond strongly to spatially structured stimuli, even when there is no net-change in the average illumination of the receptive field (Enroth-Cugell and Robson, 1966; Hochstein and Shapley, 1976; De Monasterio, 1978; Victor and Shapley, 1979; Schwartz et al., 2012) – a characteristic the LN model cannot replicate. This spatial nonlinearity is mediated via functional subunits in the receptive fields of retinal ganglion cells. These enable various specific computations that would be impossible without them, from sensitivity to fine spatial structures to various types of motion and pattern sensitivity (Ölveczky et al., 2003; Münch et al., 2009; Zhang et al., 2012; Krishnamoorthy et al., 2017; Zapp et al., 2022; Krüppel et al., 2023). Moreover, nonlinear spatial integration also plays a major role in shaping ganglion cell responses to natural stimuli (Cao et al., 2011; Turner and Rieke, 2016; Karamanlis and Gollisch, 2021). The biological correlate of receptive field subunits are the retina’s bipolar cells, which have been found to rectify their excitatory inputs to ganglion cells (Demb et al., 2001; Schwartz et al., 2012; Borghuis et al., 2013).

Given the importance of spatially nonlinear processing in the retina, understanding the underlying circuits is highly desirable. For retinal ganglion cells, extracellular recordings with multi-electrode arrays (Meister et al., 1994) allow large-scale functional characterizations that can capture the diversity of cell types (Masland, 2012; Baden et al., 2016; Grünert and Martin, 2020; Goetz et al., 2022). For bipolar cells, progress has been made on large-scale recordings using glutamate imaging (Marvin et al., 2013; Franke et al., 2017), but it is difficult to combine these techniques to obtain information about the connectivity between large populations of bipolar and ganglion cells. As an alternative, several approaches have been developed to infer subunits and thus connected bipolar cells from ganglion cell recordings alone (Freeman et al., 2015; Liu et al., 2017; Shah et al., 2020). However, these approaches are based on ganglion cell responses to fine spatiotemporal white noise stimuli, which generally evoke comparatively weak responses, making long recordings necessary. Limited recording time may thus obstruct reliable subunit identification or curtail the functional investigations of subunit layouts.

To address this issue, we introduce a novel method to identify the layout of subunits composing a ganglion cell’s receptive field that makes use of stimuli specifically targeted towards the computational characteristics of subunits and that has the potential of considerably reducing the required recording time. The method combines the functional principles underlying stimulated emission depletion (STED) microscopy (Hell and Wichmann, 1994; Hell, 2007) and tomographic imaging such as computed tomography (CT) scans (Natterer, 2001), and we term it super-resolved tomographic reconstruction (STR), accordingly. In this manuscript, we investigate the potential of STR via comprehensive modelling and electrophysiological recordings from primate retinal ganglion cells.

## Results

### Super-resolved tomographic reconstruction approach

Retinal ganglion cells often display spatially nonlinear integration of luminance signals (Enroth-Cugell and Robson, 1966). Figure 1A, B exemplifies this with responses of an On parasol cell recorded in the isolated marmoset retina. The cell responded strongly to increases of luminance during a full-field stimulus, but not to decreases of luminance (Fig 1A). On the other hand, a stimulus that involved reversals of a spatial pattern while keeping the mean luminance inside the receptive field constant also led to substantial responses (Fig 1B). This nonlinear spatial integration of luminance inside the receptive field of a ganglion cell is mediated via so-called subunits, which are believed to tile the receptive field and to correspond to bipolar cells, the source of the excitatory input to ganglion cells (Hochstein and Shapley, 1976; Victor and Shapley, 1979; Demb et al., 2001; Schwartz et al., 2012; Liu et al., 2017).

**Figure 1:**
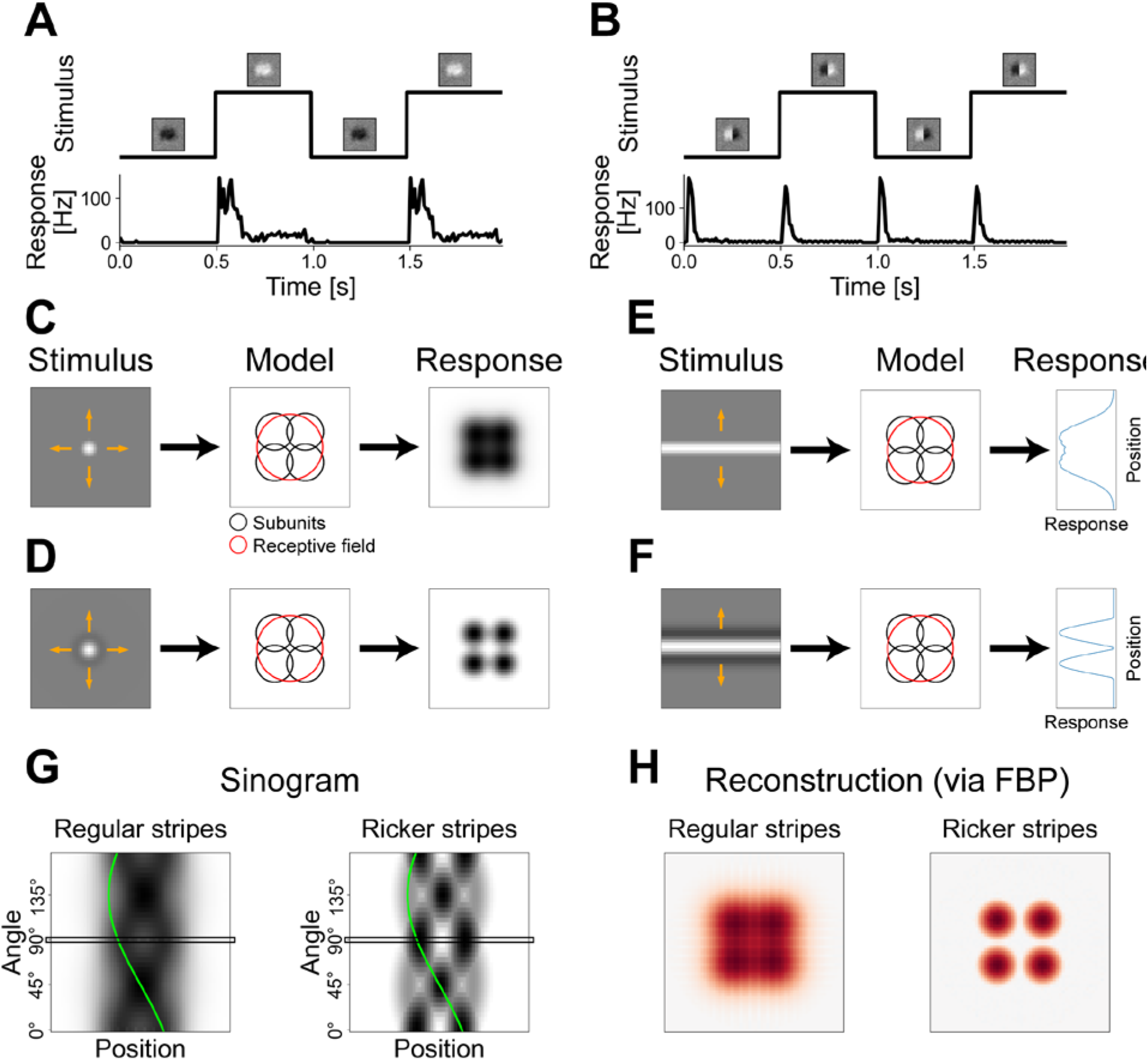
Schematic of super-resolved tomographic reconstruction (STR) method. (A) Response of a sample On parasol retinal ganglion cell to light-intensity steps without spatial structure. Top: Stimulus time course with the insets displaying a point-wise multiplication of the cell’s receptive field with the current stimulus. Bottom: Peri-stimulus time histogram (PSTH) of the cell in response. For visualization purposes, the PSTH is repeated once. (B) Same cell as in (A), but for a grating stimulus with spatial structure. (C) Schematic depiction of using a spot stimulus (left) to probe the receptive field of a model On-type ganglion cell (center). Orange arrows on top of stimulus signify the shifts of the stimulus for probing the receptive field. Black circles represent the 1.5 σ ellipses of the subunits, red circle represents the 1.5 σ ellipse of the receptive field. Response of the model (right) shows which spot positions led to a strong response (black) and which to a weak response (white). (D) Same as (C), but for a stimulus with an added dark ring around the white spot. (E) 1D probing of model responses with a horizontal stripe (left) at different vertical positions in the receptive field of the same model. The response (right) depended on the vertical position of the stripe. (F) Same as (E), but for a Ricker stripe, which has added dark sidebands adjacent to the white center stripe. (G) Sinograms of the responses of the model to the plain stripes (left) and to the Ricker stripes (right) as measured from 36 stripe angles (steps of 5°) and 60 stripe positions (steps of 2/3 pixels). Dark colors denote stronger responses. Black rectangle at 90° marks measurement shown in (E) and (F). Green line indicates sine trace of one subunit in the model’s layout. (H) Reconstructions of the sinograms in (G) using filtered back-projection (FBP). Red colors denote positive values in the reconstruction.

To develop a method for identifying the layout of subunits by recording ganglion cell responses to visual stimuli, we set up computational subunit models to simulate the response to a flashed spatial pattern. For concreteness, we focused the models on On-type ganglion cells. We modelled the nonlinear spatial integration by approximating both the subunits, i.e., bipolar cells, and the ganglion cell itself as separate LN integration stages, yielding an LNLN-like model. The first stage consisted of a linear spatial integration at the level of the subunits, which we portrayed as 2D Gaussian filters applied to the image’s Weber contrast values, followed by a rectification. At the level of the ganglion cell, the subunit outputs were then summed before a final transformation resulted in the model’s response, given as the average spike count or firing rate elicited by the flashed spatial pattern.

A simple approach to probe the spatial sensitivity profile of a ganglion cell is to flash a small spot of light (Fig 1C, left) at different locations across the cell’s receptive field. As illustrated for a schematic model with four circular subunits (Fig 1C, center), the responses of the model (Fig 1C, right) together map out its receptive field, which corresponds to the union of the subunits. Since the subunits have no gaps between each other and even overlap, individual subunits can usually not be identified by reproducing the receptive field.

To enable detection of the subunit structure inside a receptive field, our approach is to add a suppressive dark ring, or annulus, around the excitatory spot, leading to a shape that is commonly known as a Mexican hat (Fig 1D, left). By applying this stimulus, we make use of the linear luminance integration within subunits and the nonlinear integration across subunits. For instance, the average luminance of the hat is the same as the background gray (similar to the example of Fig 1B), but the model still produces responses due to its spatially nonlinear computation (Fig 1D, right). If the size of the hat is similar to the size of the subunits, an individual subunit will only be activated by the stimulus if the hat is placed close to its center. In this case, the suppressive ring of the stimulus plays a subordinate role for that subunit, due to the subunit’s greater sensitivity closer to its center, which is a reasonable assumption even if the exact Gaussian shape is a simplification. On the other hand, if the stimulus is placed at the overlap of two or more subunits, each subunit will be triggered significantly less or not at all, because the suppressive ring now strikes the more sensitive central parts of the subunits, while the excitatory white spot only hits the periphery of each of the subunits.

Comparing the response maps obtained with the simple homogeneous spot (Fig 1C) and with the Mexican-hat shaped spot (Fig 1D), similar response strengths are obtained for spots placed at the center of a subunit, because a suppressive ring does not decisively affect that subunit and the suppression of neighboring subunits evoked by the ring is rectified anyway. By contrast, for spots placed at the overlap of subunits, the suppressive ring diminishes responses, as explained above, whereas a simple spot still leads to a strong response because the partially activated subunits do not receive any suppression and combine their activation to accumulate a strong integrated response. Thus, the suppressive ring effectively leads to a spatial sharpening of the subunits, such that they can be identified more clearly as hotspots in the model’s response.

Yet, probing a receptive field with such hat stimuli would be inefficient for two reasons. Firstly, responses of a ganglion cell to an individual hat stimulus would likely be relatively weak and might not even reach spiking threshold, since only a small portion of the receptive field is activated. Secondly, the receptive field would have to be scanned point-by-point over two spatial dimensions requiring a large number of presentations at different locations.

We thus extended the idea of the spatial sharpening of subunit responses by turning towards a tomographic version of the concept. If an extensive white stripe is flashed inside the receptive field (Fig 1E, left), the response of the model will reflect the combined receptive field intensity lying within the confines of the stripe. Accordingly, if multiple positions in the receptive field are probed by such a flashed stripe, the responses will represent a projection of the receptive field along the stripe’s orientation (Fig 1E, right). Similarly, if the profile of the stripe utilizes the same shape introduced before with an excitatory center band and suppressive sidebands (Fig 1F, left), the responses will represent a projection of the sharpened subunits (Fig 1F, right). We term such a stimulus a Ricker stripe, since we apply a profile that is given by the Ricker wavelet (see Methods).

While individual subunits often cannot be identified in one such projection (in the example, both response peaks are caused by two subunits each), multiple differing projections can be measured by varying the angle of the Ricker stripe. The resulting data can be displayed in a so-called sinogram, in which each row corresponds to one projection at a fixed angle (Fig 1G). In the sinogram, every subunit leaves a sine-like trace that overlaps with the traces of other subunits for some but not all angles (green line in Fig 1G highlights the trace of a sample subunit), and their traces can thus be disentangled.

To reconstruct the subunit layout from a set of projections compiled in a sinogram, we used filtered back-projection (FBP), one of the most common techniques in the field of tomography (Brooks and Di Chiro, 1976; Willemink and Noël, 2019). The FBP reconstruction of measurements with plain white stripes (Fig 1H, left) resembles the model’s receptive field that can also be measured by probing with a plain white spot (Fig 1C). Meanwhile, the reconstruction of measurements with Ricker stripes reveals the locations of the subunits as hotspots (Fig 1H, right) similar to probing with a hat stimulus (Fig 1D). This tomographic presentation, however, has the advantage of evoking stronger responses since subunits are triggered more often and partially simultaneously, thereby reducing experiment time. We will refer to this technique as super-resolved tomographic reconstruction (STR): While the center-surround structure of the Ricker stripes sharpens subunit responses and thereby effectively super-resolves them below the scale of the subunits themselves, the presentation of stripes at varying positions and angles allows a tomographic reconstruction of subunits.

### Assessment of subunit identification with simulated data

We extended the overly simplified ganglion cell model used in Figure 1 to test STR in a more realistic setting. Firstly, we adopted a procedure to randomly generate layouts of larger numbers of subunits and also took elliptical subunits with varying degrees of overlap into account. Figure 2 shows sample layouts for 6, 10, and 14 subunits (first column). Just as before, the subunits cannot be identified from the structure of the receptive fields alone, even if noise-free high-resolution measurements are considered (Fig 2, second column). The sinograms gathered via the tomographic presentation of Ricker stripes (Fig 2, third column) show a more complex structure than in the previous, simple example of Figure 1, and individual subunit traces are progressively harder to distinguish as the model comprises more subunits. Nevertheless, the corresponding reconstructions (Fig 2, fourth column) display a clear hotspot structure with each hotspot representing the location of a subunit.

**Figure 2:**
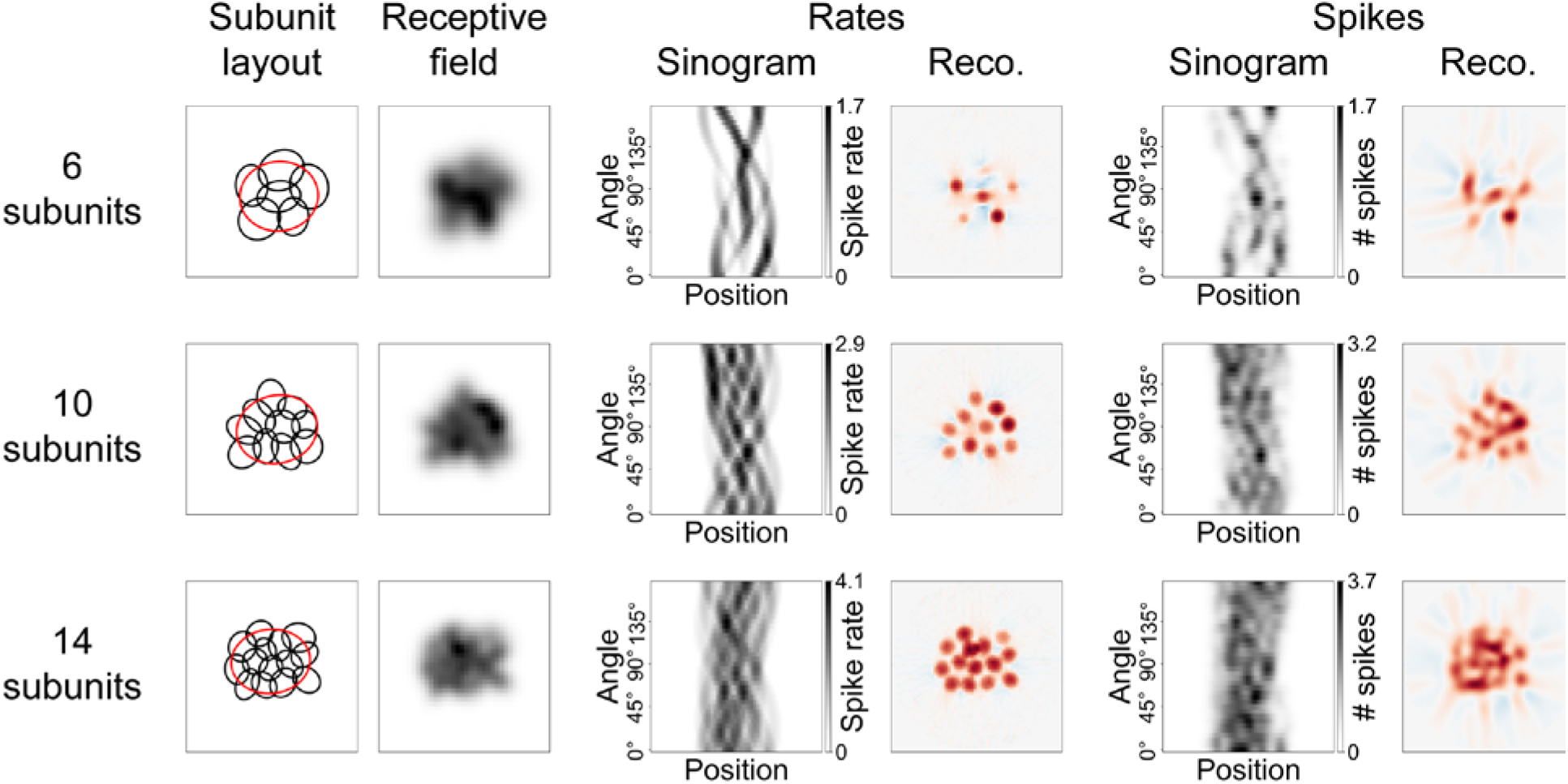
Application of STR to model simulations with realistic settings. Three sample layouts with 6 (top row), 10 (middle row), and 14 (bottom row) subunits are depicted. First column shows the subunit layouts, with black ellipses portraying the 1.5 σ ellipses of the subunits and red ellipses the 1.5 σ ellipses of the receptive fields. Second column is the receptive field. Third column contains the sinograms if spike rates, i.e., expected spike counts, are collected. Fourth column shows the reconstructions from the sinograms in the third column. Red colors denote positive values, blue colors negative values. Fifth column holds the sinograms when stochastic spike counts are measured. Each combination of the 36 stripe angles and 60 stripe positions was measured only once. Gaussian smoothing has been applied to these sinograms (described in more detail in the main text). Last column pictures reconstructions from the sinograms in the penultimate column.

Next, we extended the model to include the stochasticity of spiking responses by applying a Poisson process for spike generation in order to reflect the inherently noisy responses of real retinal ganglion cells. The Poisson process was used for simplicity, even though ganglion cell spiking is typically more regular than Poisson noise would suggest, so that this can be viewed as a conservative assessment of the effect of spiking variability. To tune the range of obtained spike counts (and thereby the noise level in the simulation), we defined how the expected spike count from the Poisson process relates to a given activation of the model. To do so, we first assumed that a full-field flash of 100% Weber contrast (i.e., “white”) would elicit an average response of 30 spikes, whereas no stimulation yielded zero spikes. For any given summed subunit signal in response to flashing a Ricker stripe, we then obtained the expected number of spikes by linear interpolation.

The sinograms in the third column in Figure 2 thus depict the expected spike count (i.e., spike rate in terms of spikes per stimulus), while the penultimate column contains the actual stochastically simulated spike count in response to one flash of the Ricker stripe for each angle and position. Although the addition of stochasticity to the model visibly affects the quality of the sinograms, the resulting reconstructions (Fig 2, last column) still feature apparent hotspot structures, although there is not always a clear one-to-one correspondence of hotspots and simulated subunits. Yet, many subunits can nonetheless easily be identified, demonstrating the potential of STR. Note that the effect of stochasticity could be reduced and reconstructions improved by averaging over multiple presentations of the same Ricker stripe, which would, on the other hand, correspond to longer recording times.

The more realistic model settings introduced above enable us to make a more meaningful assessment of the influence of certain stimulation and analysis parameters. For instance, to obtain the results presented in Figure 2, we made the following adjustments that we will also use as a default for the rest of this manuscript: We chose the width *w* of the Ricker stripe profile, defined as the distance of the two zero-crossings, i.e., the width of the white central stripe, to be *w* = 5 pixels. For comparison, in the case of models with 10 subunits, the average effective subunit diameter (diameter of a circle with the same area as the subunit ellipse) lies at 7 pixels, and the average effective receptive field diameter at just under 17 pixels. We also strengthened the sidebands of the Ricker stripes by a multiplicative surround factor *s* = 2.5 to increase their effect of sharpening the subunits. This entails that a Ricker stripe produces a net darkening, since its integral is equal to zero only if *s* = 1. Finally, sinograms obtained from models employing a spiking process were smoothed with a Gaussian of a standard deviation of *σ*_pos_ = 1.5 stripe positions and *σ*_ang_ = 1 angle step size (5°) to alleviate the influence of noise (hence the non-integer grayscale values in the penultimate column of Figure 2).

In order to assess the influence of any of these changes, we attempted to quantify the quality of a reconstruction in terms of how truthfully it reflects the subunit layout. Figure 3A illustrates the subunit layout (1.5 *σ* ellipses) and receptive field of a sample model cell that is analyzed throughout Figure 3, and Figure 3B shows the FBP reconstruction as obtained from stochastic spikes. As noted before, a successful reconstruction is characterized by the hotspots coinciding with the locations of the subunits. We therefore first located the hotspots in the reconstruction by finding all local maxima larger than 30% of the global maximum (Fig 3B, yellow markers). We discarded any hotspots that might have fallen outside a circle with 90% the diameter of the reconstructed area (Fig 3B, blue circle) to avoid artifacts, which often occurred at the edge of the reconstructed image. This procedure is deliberately simplistic since our aim here is not to present a state-of-the-art detection algorithm for hotspots, but to merely provide a means for a quantitative analysis of the reconstruction.

**Figure 3:**
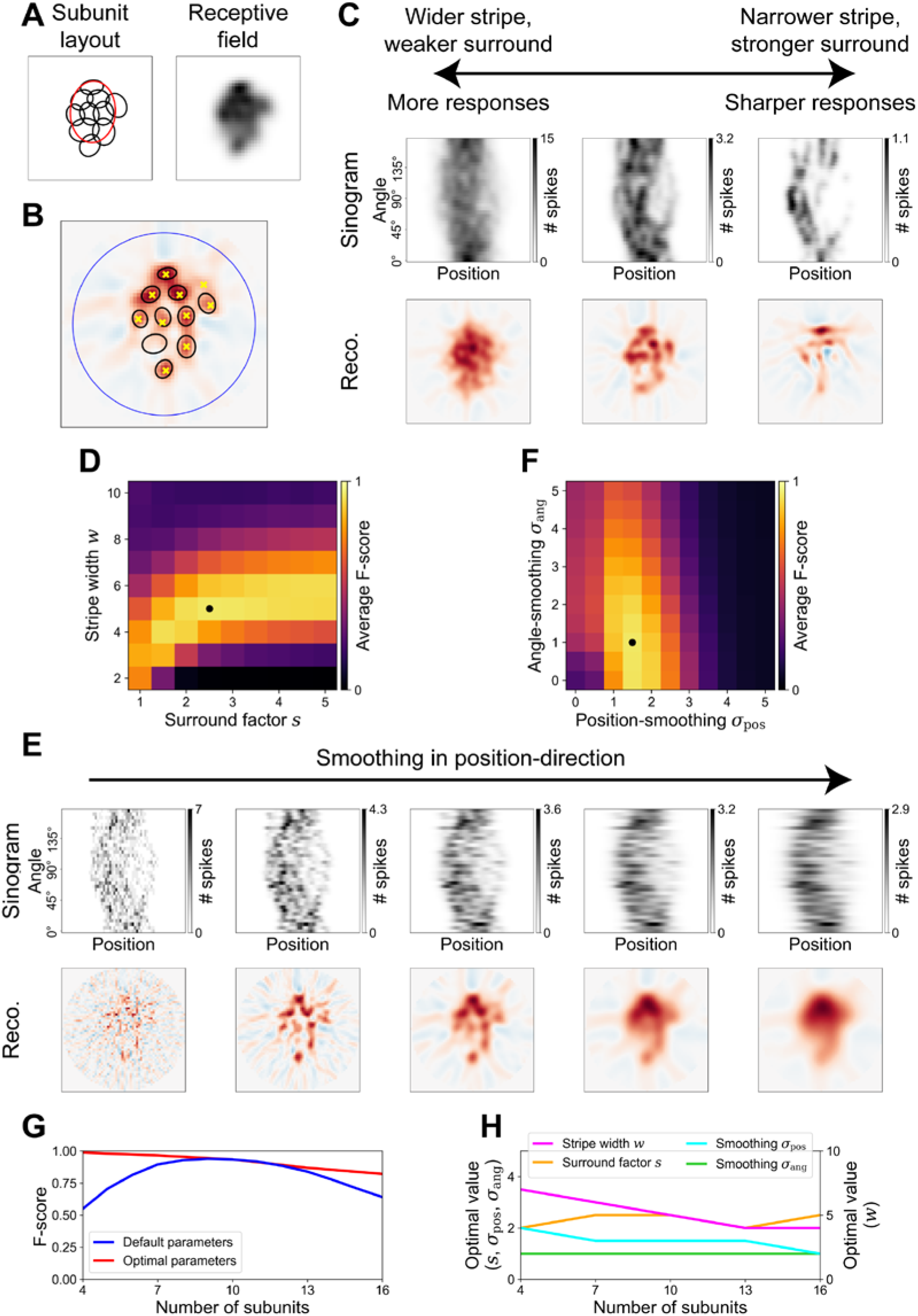
Optimal stimulus and analysis parameters. (A) Sample model layout (left) and receptive field (right) used throughout this figure (layout outlines 1.5 σ ellipses of subunits and receptive field). (B) Illustration of the detection of hotspots in a reconstruction. Background image is FBP reconstruction with red and blue colors representing positive and negative values, respectively. Large dark-blue circle depicts area in which local maxima (yellow crosses) are identified. Local maxima are compared with 0.75 σ ellipses of the underlying subunits (black) to compute an F-score. (C) Sample sinograms (top row) and corresponding reconstructions (bottom row) of measurements with varying stimulus parameters. Surround factors *s* are 1, 2, and 5 from left to right, stripe width values *w* are 10, 5, 4. (D) Search for the optimal parameters in the parameter space of surround factor *s* and stripe width *w*. Brighter colors denote better average F-score for 1000 model instantiations with ten subunits each. Optimal parameters (*s* = 2.5, *w* = 5 pixels) are marked by a black dot. (E) Influence of smoothing the sinogram in position-direction on a sample sinogram (top row) and the corresponding reconstructions (bottom row). Standard deviations *σ*_pos_ of the Gaussian filters are (from left to right) 0, 1, 2, 3, and 4 stripe positions. Smoothing in angle-direction is omitted for these plots (*σ*_ang_ = 0). (F) Like (D), but for search for optimal smoothing of the sinogram in the parameter space of standard deviations for stripe position smoothing *σ*_pos_ (optimum is 1.5 stripe positions) and stripe angle smoothing *σ*_ang_ (optimum is 1 angle step size, i.e., 5°). (G) Dependence of the average F-score (over 1000 model instantiations) on the number of subunits of the models. Blue curve characterizes dependence for the default parameters (optimal for 10 subunits) identified in (D) and (F), red curve for parameters optimized for 4, 7, 10, 13, and 16 subunits, respectively. (H) Trends portray how the optimal parameter values depend on the number of subunits. Optimal parameters were manually determined for 4, 7, 10, 13, and 16 subunits, and reflect those used for generating the red curve in (G).

After finding the hotspots, we determined which of them fell inside the 0.75 *σ* ellipse of a subunit (Fig 3B, black ellipses; double hits counted as one hit and one miss) and calculated the F-score, which is a combined measure of precision and sensitivity (see Methods). The F-score ranges from 0 to 1, with values close to 0 indicating that few subunits had been detected via hotspots (low sensitivity) and/or that there were many hotspots that did not correspond to subunits (low precision), and 1 indicating that hotspots and subunits matched perfectly. In the example of Figure 3B, nine of the ten detected hotspots lay in the 0.75 *σ* ellipse of a subunit, and one subunit was not detected, leading to an F-score of 0.9. Since the noise of the spike generation is an integral part of the challenge of choosing good stimulus and analysis parameters, all considerations made below, including the calculation of F-scores, assume models with a stochastic spike generation process as in Fig 3B.

With this evaluation system in place, we first examined how the quality of the reconstruction depends on the characteristics of the Ricker stripes, i.e., their width *w* and sideband strength *s*, while fixing the sinogram smoothing parameters *σ*_pos_ and *σ*_ang_ at the default. As expected, by increasing the width of the stripes and decreasing the strength of the suppressive sidebands, the model produced a larger number of spikes in response (Fig 3C, left). While this means that the sinogram is less noisy, it also entails that the reconstruction is more representative of the receptive field instead of the individual subunits. On the other hand, a narrow stripe with strongly suppressive sidebands, supposed to sharpen the subunit responses substantially, results in responses too weak for FBP to reveal many subunits (Fig 3C, right). We thus probed the parameter space of Ricker stripe width *w* and surround factor *s* to discover if a proper balance can be found. Figure 3D shows the average F-score STR achieved for layouts of ten subunits, and, indeed, certain combinations of the two parameters led to respectable F-scores. The best F-score of 0.93 was reached using the previously introduced combination of the stripe width *w* = 5 pixels and the surround factor *s* = 2.5 (Fig 3D, black dot).

Similarly, we investigated the effectiveness of smoothing the sinogram in order to overcome the issue of noise. Figure 3E exemplifies the influence smoothing in the stripe position-direction if omitting any angle-smoothing and demonstrates that the noise introduced with the spike generation process greatly impairs the FBP reconstruction if not counteracted (leftmost example involves no smoothing at all). On the other end of the spectrum, too much smoothing naturally eliminates all finer structures and the reconstruction simply reflects the receptive field. Again, a balance must be found and we computed the average F-score of layouts with ten subunits to identify this balance (Fig 3F). Doing so, we determined the optimal standard deviation for Gaussian smoothing to be *σ*_pos_ = 1.5 stripe positions and *σ*_ang_ = 1 angle step size (5°) – the values that we adopted as our default.

While we here determined the optimal stimulus and analysis parameters for layouts of ten subunits, these parameters also worked well for more or fewer subunits. Figure 3G shows that, for a range of different subunit numbers, subunit detection was similarly effective with the default parameters (blue line) as with parameters optimized for each specific number of subunits (red line; obtained by alternatingly optimizing the pair *w* and *s* and the pair *σ*_pos_ and *σ*_ang_ as in Fig 3D and F). In addition, this analysis also revealed that both small and large numbers of subunits could be identified well with the right parameters, but layouts with many subunits were somewhat harder to uncover. This is likely an effect not of the increased number of subunits itself, but rather of the decreased effective resolution in our simulations when a larger number of subunits was used. Since our layouts were scaled to a similar receptive field size, there were fewer pixels and stripe positions available per subunit in layouts with more subunits. This naturally complicates distinguishing them in the reconstruction. Regardless, certain trends of the optimal parameters are apparent for varying numbers (or sizes) of subunits (Fig 3H). First, the optimal values for both the surround factor *s* and the angle-smoothing *σ*_ang_ remained largely unaffected by changing the number of subunits. Thus, standard values for these parameters can be applied in the analysis regardless of how many subunits are expected. For both the stripe width *w* and the smoothing in position-direction *σ*_pos_, on the other hand, the optimal values decreased for higher numbers of subunits. This is expected, as in our simulations more subunits correspond to smaller subunits, suggesting an analysis at finer spatial scales and adjusting the stripe width to the approximate expected size of subunits.

### Robustness of method

So far, the considered subunit models were all based on the same general structure with fixed characteristics, like the subunit nonlinearity and the shape and overlap of subunits, and only varied in the specific subunit layouts. However, different ganglion cell types have different functional properties, which can also vary depending on, e.g., retinal location (Bleckert et al., 2014; Warwick et al., 2018), and many specifics are not known a priori. Identifying subunits with STR, however, is robust against variations in subunit signaling, as revealed by changing various model components.

For example, we increased the overlap of the subunits leading to receptive fields with even less discernible structure (Fig 4, top row, left plots show sample layout and corresponding receptive field). Nevertheless, noise-free sinograms comprised of firing-rate responses appear not visibly inferior to those from the default model composition, and reconstructions calculated from these sinograms still clearly represent the subunit layout (Fig 4, top, center). When a spiking process is included in the model, sinograms and reconstructions also demonstrate decent quality (Fig 4, top, right), with the average F-score decreasing somewhat to 0.83. Much of this loss can be recovered by increasing the stripe width from *w* = 5 to *w* = 6 pixels and the surround factor from *s* = 2.5 to *s* = 3. This is because the increased overlap of subunits is counteracted by a stronger response sharpening, which results from the larger surround factor. In addition, the increased overlap in our simulations goes along with a larger size of each subunit compared to previous layouts, which makes the wider stripes more suitable. Together, this leads to an F-score similar to one for the default scenario, which might be unexpected, since an increased overlap should make distinguishing the subunits more difficult. However, as we have noted above when investigating the influence of the number of subunits, the effective resolution per subunit is an important factor governing the quality of the reconstruction, and is improved with the larger subunits.

**Figure 4:**
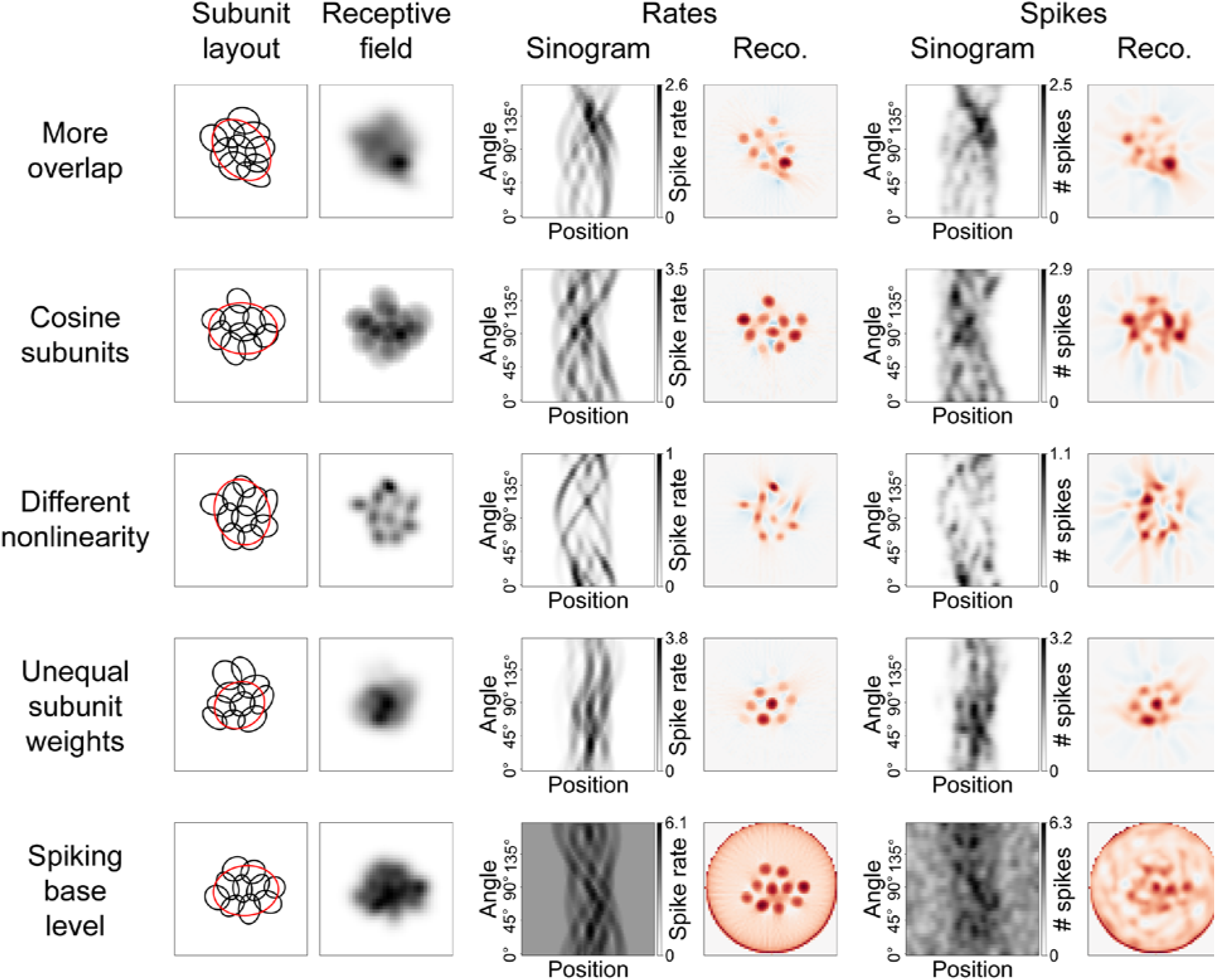
Robustness of STR to model variations. Each row demonstrates the effect on STR of one variation of the model via a sample simulation. Layout of the rows is the same as in Fig 2. Top row shows a model with increased subunit overlap (see Methods for details) apparent from the 1.5 σ subunit ellipses. In the second row, the Gaussian-shaped subunits were replaced with subunits of a cosine profile. For comparability, the ellipses in the subunit layout depiction are 1.5 σ ellipses of Gaussians fitted to the cosine subunits. Third row contains a replacement of the rectified-linear nonlinearity with a rectified-quadratic nonlinearity. Weights of the subunits in the fourth row were not all equal as in the default model, but chosen according to a large spatial Gaussian. In this example, the strongest subunit weight was roughly eight times that of the weakest weight. In the bottom row, a base activity of three expected spikes was added to all responses.

Furthermore, this also increases the size of the 0.75 *σ* ellipses used to determine if a hotspot corresponds to a subunit, thereby leading to a potentially better F-score even if the exact same reconstruction would be obtained. Altogether, we conclude that an increased subunit overlap is not per se detrimental to STR.

The exact shape of the subunits is also not a sensitive factor. We have replaced the Gaussian subunits of the standard model with subunits, whose cross-sections reflect the positive part of a cosine curve between zero-crossings (Fig 4, second row; for comparability, subunit ellipses depict the 1.5 *σ* ellipses of Gaussians fitted to the cosine subunits). Again, the reconstructions clearly exhibit a hotspot structure that reflects the subunit layout and the average F-score increased marginally to 0.94.

Other changes to the model’s properties, like modifying the subunit nonlinearities, can have stronger effects. We replaced the default rectified-linear transformation of subunit signals with a rectifying nonlinearity that additionally squared all positive values (Fig 4, third row). Doing so leads to the subunits being directly apparent in the receptive field, because weak excitation of multiple subunits at their overlap evokes a weaker model response than strong excitation of a single subunit at its center due to the squaring of subunit signals. Note, however, that Fig 4 shows a noise-free high-resolution receptive field measurement that is virtually impossible to achieve in a real experiment, and a more realistic spike-triggered average (STA) under spatiotemporal white noise would likely not reveal any subunits. While the reconstruction of noise-free responses to Ricker stripes still flawlessly represents the subunits, the reconstruction obtained from stochastic spiking responses was certainly undermined by the nonlinearity change, with the average F-score decreasing to 0.76. This performance loss seems to stem from the noticeably weaker responses to the Ricker stripes (cf. the ticks at the sinograms’ grayscale bars) and thus inferior signal-to-noise ratio. This, in turn, is a result of our choice to set the reference point for the spike count at the maximum response, that is, the response to a full-field white flash. For Ricker stripe stimuli, which activate the model less strongly than the full-field flash, the quadratic nonlinearity therefore weakens the responses compared to the rectified-linear nonlinearity. Indeed, the F-score can be improved up to 0.87 by making the Ricker stripes wider from *w* = 5 to *w* = 6 pixels, which, as noted in Figure 3, increases the response strength.

In the simulations, we had so far assumed all subunits to contribute with an equal weight to the response of the model, but subunits might realistically contribute differentially, with subunits farther from the cell’s center potentially having a weaker connection to it. We modelled this hypothesis by choosing subunit weights according to a 2D spatial Gaussian that prefers subunits close to the center. Consequently, subunits in the periphery of the receptive field played a minor role in activating the modeled ganglion cell, a fact demonstrated by the top two subunits in the sample layout in Figure 4 (fourth row) being almost irrelevant for its receptive field. While more central subunits are still well represented in the reconstructions, these outer subunits are difficult to recognize, thus reducing the average F-score to 0.76. On the other hand, the significance of the F-score is limited in this case, because it values all subunits equally, which does not reflect their true contribution to the model’s responses.

As a final variation of the standard model, we added spontaneous activity to the measurements (Fig 4, bottom row), which we implemented by increasing the expected number of spikes for the Poisson process by 3 irrespective of the stimulus. This is a considerable level of background activity, as it is similar to the maximum systematic response modulation evoked by our stimuli. One effect on the FBP reconstruction is a characteristic artifact ring at the periphery of the reconstructed image. This in itself does not strongly compromise the subunit reconstruction, as is apparent from the high quality of the noise-free reconstruction. Yet, for the automated hotspot detection, we had to confine potential hotspot locations to a circle with a diameter of 90% of the reconstruction region. More importantly, however, the overall noise in the reconstruction is strongly increased by adding spontaneous activity, leading to a decrease in the average F-score to 0.59. The F-score could be improved to 0.68 by increasing the standard deviations of the Gaussian smoothing from *σ*_pos_ = 1.5 to *σ*_pos_ = 2 stripe positions and from *σ*_ang_ = 1 to *σ*_ang_ = 1.5 angle step sizes (7.5°), thereby counterbalancing the noise to some degree. Yet, substantial levels of background noise can evidently have a significant influence on the quality of the FBP reconstruction.

While FBP is a simple and easy-to-use algorithm to reconstruct the subunit layout, it is not designed for our specific reconstruction problem. We therefore considered how the specifics of FBP may limit its suitability for subunit detection. Aside from the susceptibility to noise in the measurements, as discussed above, a particular caveat concerns the reconstruction of elliptical subunits. Indeed, the circular symmetry of subunits that we had assumed in Figure 1 is clearly an abstraction, and elliptical shapes with different levels of eccentricity are rather the norm in the retina (Liu et al., 2017; Shah et al., 2020).

For our stimulation with Ricker stripes, this means that the mean luminance inside an elliptical subunit, even if the stripe precisely hits its center, can still depend heavily on the stripe’s orientation. If the stripe is oriented perpendicular to the major axis of the subunit (Fig 5A, left), the suppressive sidebands will influence the subunit much more than if the stripe was oriented parallel to it (Fig 5A, right). Consequently, the response elicited by that subunit depends drastically on the angle of the stripe and might even be completely suppressed for some angles (Fig 5B). In X-ray tomography, this phenomenon would correspond to strongly anisotropic absorption, which FBP does not take into account. Instead, the reconstructed subunit is smeared out along its major axis and negative troughs are reconstructed adjacent to it (Fig 5C, red denotes positive values, blue negative values). While the elliptical subunit can still readily be identified in this simple example consisting of just the one subunit, in more complex layouts these effects can overlap and undermine the quality of the FBP reconstruction. The extent of this effect also depends on the subunit nonlinearity – the rectifying and squaring nonlinearity mentioned before, e.g., will magnify this issue. Nevertheless, as demonstrated before, FBP reconstructions still reproduce the locations of many subunits, but by taking the effects described here into account, an alternative reconstruction method might help locate subunits more consistently, reduce trough effects in the reconstruction, and determine the subunit shape more accurately.

**Figure 5:**
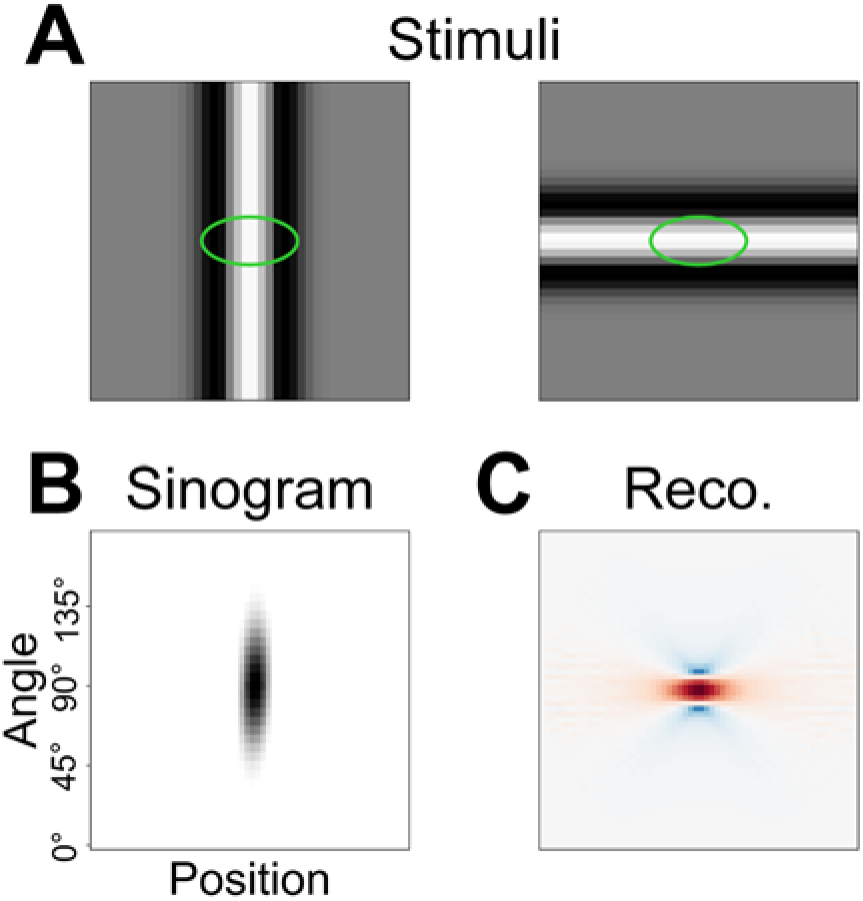
Shortcomings of FBP as a reconstruction method for subunit layouts. (A) Sample stimuli of a vertical (left) and a horizontal (right) Ricker stripe hitting the center of an exemplary elliptical subunit (here in green). (B) Rate responses of a model consisting of only that one subunit depicted as a sinogram. (C) Resulting reconstruction via FBP from the sinogram in (B). Red colors denote positive values, blue colors negative values.

### Experimental test

In order to experimentally test our method, we performed an electrophysiological recording of ganglion cells in an isolated marmoset retina using a multielectrode array while projecting light stimuli onto the retinal photoreceptors. We identified On and Off parasol cells, which we expected to exhibit the most pronounced subunit structure due to their spatially nonlinear computation, from their responses to a spatiotemporal white noise stimulus. Both cell types displayed a fast biphasic filter, tiling of visual space by their receptive fields, and consistent autocorrelation functions and output nonlinearities (Fig 6A). Off parasol cells had an average effective receptive field diameter of 111 ± 9 µm (mean ± standard deviation) and On parasol cells of 138 ± 11 µm, both values in line with dendritic field sizes in the marmoset peripheral retina (Masri et al., 2019).

**Figure 6:**
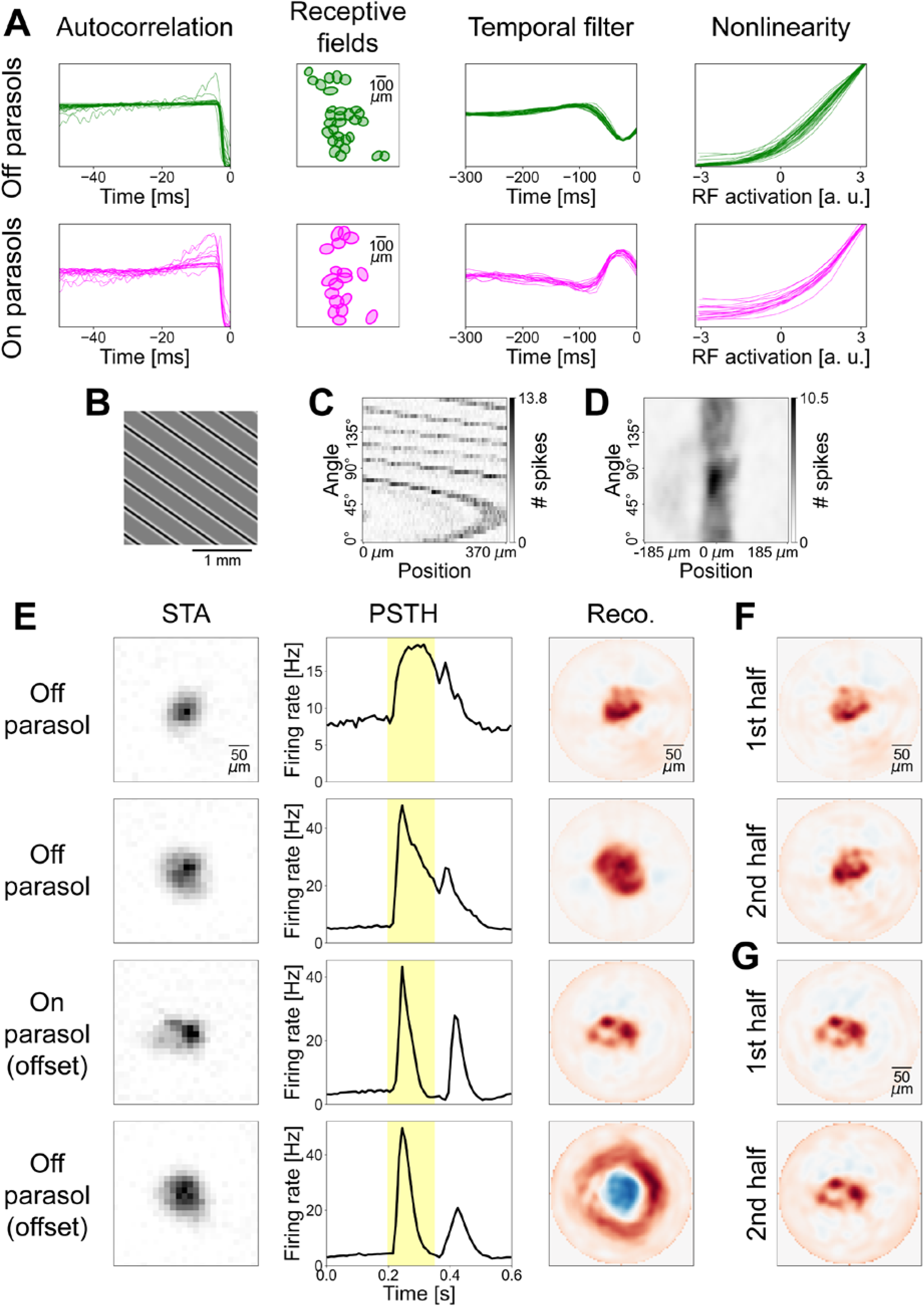
Experimental application of STR. (A) Autocorrelation functions, receptive fields (RFs) displayed as 1.5 σ ellipses of Gaussian fits (one distant cell not included), temporal STAs (normalized to unit Euclidean norm), and nonlinearities (scaled to equal maximum) of all identified Off (top) and On (bottom) parasol cells. (B) Excerpt of a sample stimulus projected onto the retina during a flash. (C) Unprocessed sinogram of a sample Off parasol cell. (D) Same sinogram as in (C), but processed by correcting for the receptive field position and applying a Gaussian filter. (E) Overview of the results of four sample cells. Left column depicts spatial STAs, middle column illustrates PSTHs (yellow background designates presentation of the Ricker stripes), right column shows reconstructions of the processed sinograms via FBP. Red colors in reconstructions denote positive values, blue colors denote negative values. Spatial scales of STAs and reconstructions are equal, but reconstruction has higher resolution. PSTHs were computed irrespective of the angle and position of the stripes with a bin size of 10 ms. Sample cell in top row is same cell as in (C) and (D). Bottom two rows contain the results of an analysis of the offset responses of cells. (F) Reconstructions of the sample Off parasol cell from the top row of (E) from separate analyses of the first and second halves of the measurement. (G) Same as (F) but for the On parasol cell from (E).

For the experiments, we adjusted the Ricker stripe stimulus in two major ways: Firstly, we decided to mainly target Off parasol cells, which, for the macaque retina, had been shown to have stronger spatial nonlinearities in the receptive field than their On-type counterparts (Turner and Rieke, 2016; Yu et al., 2022). We therefore flipped the polarity of the Ricker stripes, i.e. now having a black center and white sidebands, which act excitatory and suppressive, respectively, for Off cells. Second, to be able to probe a larger portion of the tissue simultaneously, we employed multiple parallel Ricker stripes (Figure 6B shows an excerpt of the screen). The stripes had a safety distance of 375 µm between them to ensure that only one stripe hits a receptive field for any given stimulus presentation. The Ricker stripes were flashed for 153 ms separated by 447 ms of full-field gray, and all spikes occurring during the flash of the stripes were used to compose the sinograms. Our choice of stimulus parameters was guided by our simulations with some adjustments. In the simulations, we had used a simulated area of 40 by 40 pixels, with about 20 by 20 pixels being occupied by the receptive field of the model.

In the experiments, the effective receptive field diameters of Off and On parasol cells corresponded to 15 and 18 pixels, respectively, such that parameter values are roughly comparable. We decided to use a stripe width *w* = 6 pixels and a surround factor *s* = 1.5. Reducing the surround factor was motivated by our expectation of incomplete rectification by the subunit nonlinearity, resulting in a component of linear spatial integration, which could weaken ganglion cell responses and decrease the signal-to-noise ratio. Similarly, we chose the slightly larger stripe width to make sure that cells with a supposedly easier-to-reconstruct small number of subunits were stimulated appropriately. The number of stripe positions (75, covering a distance of 50 pixels in steps of 2/3 of a pixel) and the number of stripe angles (36) matched the simulations. In the experiments, each combination of position and angle was presented three times to yield an averaged response.

Measured sinograms (Fig 6C) showed a strong overall curvature, even extending beyond the edges of the sinogram, owing to the receptive fields – in contrast to the setting in the simulations – not being centered on the screen. Note that the sinograms here are cyclical, because each stimulus consists of multiple Ricker stripes. That is, when the overall structure in the sinogram crosses the edge, this corresponds to a transition from one specific stripe hitting the receptive field at the farthest shift in one direction, to the neighboring stripe hitting the receptive field at the farthest shift in the other direction. This effect can easily be compensated for, however, by redefining the zero-position in the sinogram. For a given angle, we set the zero position to the stripe position that was closest to the receptive field center known from the white noise stimulus. Figure 6D shows such a corrected sinogram, including a Gaussian smoothing with a standard deviation of *σ*_pos_ = 1.5 stripe positions and *σ*_ang_ = 1 angle step size applied. This makes the sinogram similar in appearance to the smoothed sinograms in the simulations analyzed above, which had led to successful detection of the model’s subunits.

STAs of Off and On parasol cells computed from an hour-long white-noise stimulus with high spatial resolution displayed generally little structure inside the receptive fields (Fig 6E, left column shows four sample cells). The average peri-stimulus time histograms (PSTHs) for the tomographic stimulus, calculated by averaging over all stripe positions and angles, demonstrate a strong response to the onset of the Ricker stripe flash and also a noticeable response to its offset for most cells (Fig 6E, middle column, yellow background marks duration of stripe flash). The upper two sample reconstructions in Figure 6E (right column) were calculated for two Off parasol cells from the responses that occurred during the stripe flashes. Hotspots are less clearly delineated as in the simulations, but can generally still be recognized.

In addition to the strong response to the onset of the Ricker stripes, the offset response can also be analyzed. This is most informative for On parasol cells, where an analysis of the offset response via the same methods described above for the onset can also reveal a hotspot structure in the reconstruction that is not apparent from the STA (Fig 6E, third row). The sample On parasol cell in Figure 6E is the best example in our dataset and shows four clearly distinct hotspots suggesting that the receptive field of that cell is composed of four subunits. Consequently, while the stimulus we used in the experiments was targeted towards Off cells, the receptive field substructure of cells of the opposite polarity can still be examined by focusing on the offset response. Finally, we also inspected the offset response of Off parasol cells, which exposed a center-surround structure in the reconstruction, but little substructure. Since the average luminance of the Ricker stripes is dominated by the sidebands (due to the surround factor *s* > 1) and spatial integration in the surround of ganglion cells is not expected to be strongly nonlinear on this scale (Takeshita and Gollisch, 2014) the surrounding ring in the reconstruction might be caused by a detection of the offset of the suppressive sidebands by the receptive field center, rather than being a response of the surround itself.

Whether the hotspot structure in the reconstructions actually corresponds to subunits and potentially bipolar cells or is just a result of the noise of the measurement is not immediately clear, as we do not have a ground truth available for the experimental data. Thus, to probe the reliability of the recovered substructure in the receptive field, we split the measurement of the tomographic stimulus into a first and second half and analyzed them separately for comparison. The resulting reconstructions demonstrate that similar receptive field substructure may result from independent data sections (Fig 6F, cell shown is top Off parasol from Fig 6E, and Fig 6G, sample cell is On parasol from Fig 6E), corroborating the biological origin of the derived structures. This was not the case for all cells in our dataset, however, as other sample cells could show misaligned hotspots from the two halves of the data or unclear structure. This could suggest that the reconstructions were governed by noise and that a reliable subunit structure may not be discernible with STR in these cells. On the other hand, responses of these cells might not have been sufficiently stable over the recording duration or using only half of the data, where some stimuli were presented only once, others twice, could be insufficient to get reliable reconstructions. Thus, while the analysis of the recorded data indicates the feasibility of the approach for real ganglion cells, more systematic experimental explorations will be required in the future.

## Discussion

Spatially nonlinear integration of luminance signals inside the receptive fields of ganglion cells is mediated via subunits, which are thought to correspond to the retina’s bipolar cells (Demb et al., 2001). Inferring the subunit layout from electrophysiological measurements of a ganglion cell promises a new avenue towards understanding the functional properties of the retina’s circuitry. Several studies have proposed methods for subunit inference (Freeman et al., 2015; Liu et al., 2017; Shah et al., 2020), but these typically require long recordings with finely structured stimuli that are inefficient in driving responses of ganglion cells. This makes it difficult, for example, to retain sufficient experiment time for studying the uncovered structure in more detail and relate it to functional analyses of stimulus encoding. We here introduced the method of super-resolved tomographic reconstruction (STR) that combines concepts underlying STED microscopy and tomography to identify subunits from retinal ganglion cell recordings. STR consists of flashing Ricker stripes – named after the Ricker wavelet that describes their center-surround profile – with varying angles at different positions in the receptive field (Fig 1).

The evoked ganglion cell responses can be arranged in a sinogram and the subunit layout can be reconstructed from that sinogram, for example by applying filtered back-projection (FBP). Simulations demonstrated that hotspots in the FBP-based reconstruction reliably corresponded to subunits (Fig 2). To optimize performance, especially when data are noisy, stimulus and analysis parameters can easily be tuned according to the response properties of the investigated neurons (Fig 3). Yet, with fixed parameters in an appropriate range, the method proved to be robust against varying the specifics of the subunit model (Fig 4), even though distortions of the reconstructed subunit layout can occur for non-circular subunits when employing FBP as a reconstruction algorithm (Fig 5). Application to recordings of parasol ganglion cells in the primate retina indicated the experimental feasibility of the approach, although further experimental explorations will be required to evaluate the reliability of the results (Fig 6).

### Relation to STED microscopy and tomography

Our super-resolution approach is conceptually related to stimulated emission depletion (STED) microscopy (Hell and Wichmann, 1994; Hell, 2007). In regular confocal fluorescence microscopy, a specimen containing fluorescent molecules is scanned with an excitatory spot of light. The resolution of this microscopy technique is determined by the size of the area that emits fluorescent light, which, in turn, is given by the size of the excitatory spot of light. Due to Abbe’s diffraction limit, however, a spot of light cannot be focused to an arbitrarily small size, thereby limiting the resolution. STED microscopy therefore introduces a second source of light, shaped like a ring, which depletes fluorescence. Any remaining fluorescent light emission is thus confined to areas that were covered by the excitation but not depletion light. Consequently, the area emitting fluorescent light shrinks, thereby improving the resolution compared to confocal fluorescence microscopy beyond Abbe’s diffraction limit.

In the context of subunit identification, the spatial extent of the subunits takes the role of Abbe’s diffraction limit. Pairs of subunits that overlap are difficult to separate by spots of light alone, just like pairs of fluorescent molecules with a distance less than the diffraction limit cannot be excited separately. However, the addition of the suppressive ring around the spot of light shrinks the area in which subunits are responsive to the stimulus. A subunit not quite centered with regards to the stimulus is excited by the central spot, but simultaneously “depleted”, i.e., suppressed, by the ring of opposing contrast, just like the fluorescence signal from an off-center molecule in STED microscopy is suppressed by the depletion ring surrounding the center of the excitation. Like in STED microscope, this effect here leads to a super-resolution of the subunit layout beyond what simple spot-like stimulation would suggest.

As a second concept, we introduced a tomographic variation of the super-resolution stimulus in order to evoke stronger responses and sample the receptive field more efficiently. Bars of light have long been used to qualitatively study receptive field properties of visual neurons (Hubel and Wiesel, 1959, 1962), and Sun and Bonds (1994) introduced the application of filtered back-projection to quantitatively determine the receptive fields of cells in the cat lateral geniculate nucleus (LGN). Since then, this approach has been employed to locate receptive fields in various systems including the goldfish, zebrafish, and mouse retina (Johnston et al., 2014; Eiber et al., 2021), primate LGN, V1, and V2 (Pipa et al., 2012; Fiorani et al., 2014; Eiber et al., 2021), and even fMRI of human visual cortex (Greene et al., 2014). Previous studies of retinal ganglion cells have shown that the tomographic analysis scheme can markedly accelerate receptive field estimations (Johnston et al., 2014; Eiber et al., 2021), compared to stimulation with individual spots of light or spatiotemporal white noise. We believe the same effect to apply in our tomographic approach with super-resolution stimuli for subunit identification, as Ricker stripes constitute a much more potent stimulus for driving ganglion cells than small spots with suppressive rings or spatiotemporal white noise at the required high spatial resolution.

Nevertheless, differences between the ganglion cell system investigated here and X-ray tomography may complicate the analysis of the sinograms and limit the applicability of FBP. The FBP algorithm provides a discrete approximation of the inverse Radon transform (Natterer, 2001). In the context of X-rays, the Radon transform describes the absorption of the beams travelling through an object. In more general words, it calculates projections of an object in various directions. If the stimulus used here was an infinitesimally narrow bar without suppressive sidebands, the Radon transform would perfectly describe the responses of our standard ganglion cell model. However, since the Ricker stripes have a finite width and the suppressive sidebands trigger the nonlinearities in the system, the Radon transform reflects only an approximate description of the system. Consequently, deviations occur in the inverse Radon transform, e.g., when the nonlinearity of an elliptical subunit is triggered differentially for different angles (Fig 5). Thus, improvements to the reconstruction of the subunit layout could come from replacing or amending FBP by an appropriate reconstruction algorithm that does not rely on the inverse Radon transform. Iterative reconstruction methods or deep learning approaches potentially combined with LNLN models to replace the Radon transform and with appropriate regularization schemes are promising starting points (Natterer, 2001; Nuyts et al., 2013; Adler and Öktem, 2018; Maier et al., 2019; Wang et al., 2020).

### Identification of subunits from reconstructions

The main objective of the present work has been to obtain reconstructions that can be viewed as a representation of the subunit layout within the receptive field of a retinal ganglion cell. In the idealized scenario of simulated, noise-free ganglion cell responses, the individual subunits are easily identifiable as hotspots in the reconstructed image. When noise or other complications were added to the simulations or when experimental data were considered, the correspondence of hotspots and subunits became less clear. For simplicity, we evaluated the reconstructions by detecting hotspots as local maxima, but future applications may take a more sophisticated approach, such as fitting a mixture of Gaussians to the reconstruction or employing advanced blob-detection algorithms (Kong et al., 2013). Clearly, however, the success of any subunit identification technique will depend on how distinct and reliable the hotspots in the reconstructed images are.

Regarding our experimental results, the observation of partially differing hotspot locations obtained from the first versus second half of the recording (Fig 6F, G) exemplifies the limits of our approach. Multiple causes for the observed differences are conceivable and most of them could be remedied in future experiments.

For example, the recording quality of the analyzed experiment may not have been sufficient. In particular, the ganglion cells measured here might not have maintained sufficiently stable responses during the recording to warrant a comparison of responses separated by about 45 minutes. Electrophysiological measurements from isolated retinas can decrease in quality over time with cells often responding more sluggish later on, which can have an effect on their response accuracy to the fine Ricker stripes. In this case, the recording’s first half (or first trial) might in fact have yielded a good correspondence of hotspots and subunits, but without knowledge of the ground truth this can hardly be determined. Simultaneous recordings of connected bipolar cells, as previously performed in the salamander retina by Liu et al. (2017), might help assess whether identified subunits do indeed represent bipolar cell receptive fields, but such experiments have been difficult, in particular in the mammalian retina.

Another potential source of hotspot and subunit discrepancy might lie in our choice of stimulus parameters. We took a conservative approach, opting for somewhat wider Ricker stripes with weaker sidebands than our simulations suggested, in order to ensure adequate response strength. That goal was easily met, but at the same time there appears to be too little sharpening of subunits, judging from the lack of distinctly separated hotspots in experimental compared to simulated reconstructions. In future applications, a general line of thought to follow when deciding on the parameters of the Ricker stripes would be to choose their width *w* to be similar to or slightly smaller than the expected diameter of subunits and increase the sideband strength *s* as much as the evoked responses permit to sharpen the contributions of individual subunits.

Furthermore, we decided to target our stimulation towards Off parasol cells by using Ricker stripes with dark centers and bright sidebands. This was motivated by previous observations that Off parasol cells receive more strongly rectified input signals than On parasol cells in the macaque retina (Turner and Rieke, 2016; Yu et al., 2022). Here, we observed that offset responses of On parasol cells are not inferior for our analysis to onset responses of Off parasol cells, which is in line with previous findings about spatially nonlinear responses of parasol cells to grating on- and offsets (Yu et al., 2022). Alternatively, it could suggest that marmoset On parasol cell measurements are more suitable for our method than expected, in line with recent observations that in the marmoset retina, On parasol cells may display particularly strong spatial nonlinearities (Karamanlis et al., 2023).

Other limits of subunit identification from experimental data may also stem from the investigated retinal circuitry itself. While On and Off parasol ganglion cells in the primate retina are often considered to receive most of their excitatory synaptic input from a single type of bipolar cells each, namely diffuse bipolar cells DB4 and DB3a, respectively, other bipolar cell types may also contribute substantially (Jacoby et al., 2000; Calkins and Sterling, 2007; Tsukamoto and Omi, 2015, 2016; Masri et al., 2016). If these create multiple superimposed subunit layouts, individual subunits would be difficult to identify from the reconstruction. Bipolar cells are also connected via gap junctions (Jacoby and Marshak, 2000), which support, e.g., motion sensitivity (Manookin et al., 2018), and these couplings could influence spatial stimulus integration.

Additional complications in the experimental data, compared to the simulations, may arise from the receptive field surround of bipolar cells (Dacey et al., 2000; Thoreson and Mangel, 2012) and from interactions with inhibitory amacrine cells. The center-surround structure, however, would be in line with the profile of the Ricker stripes and should thus rather add to the sharpening of responses when the stripe is aligned with the center of a subunit. More problematic could be the influence of amacrine cells. A multitude of different amacrine cell types are known (Yan et al., 2020), but their functions remain largely unclear (Franke and Baden, 2017). One hypothesis is a linearization of the nonlinear bipolar-to-ganglion cell synapses (Werblin, 2010; Yu et al., 2022). Such effects could hamper the success of STR, since it relies heavily on the rectification of excitatory subunit signals. Pharmacological intervention to block inhibition could be a way to better isolate the feed-forward signal processing structure investigated here.

### Implications for circuit analysis

Despite the caveats regarding the correspondence of hotspots in the reconstruction of experimental data to subunits, STR has the potential to identify subunits in a time-efficient manner. Our simulations suggest that a single trial of every stripe position and angle may be sufficient, and we partly observed significant hotspot structures in FBPs from similarly short experimental measurements (Fig 6G). The potential to acquire sufficient data with such short measurements is due to the Ricker stripes serving as a more potent stimulus for driving ganglion cell responses than, e.g., fine spatiotemporal white noise. In addition, no computationally intensive post-processing is required, such that accurate reconstructions of subunit locations could be available within just 30 minutes of experiment time, which is significantly faster than existing methods (Freeman et al., 2015; Liu et al., 2017; Shah et al., 2020). A further reduction might be achieved by optimizing the presentation times of the Ricker stripes and reducing the number of probed angles or positions, which could become feasible when more advanced reconstruction techniques are employed to deal with the reduced data.

Such a fast subunit identification leaves ample recording time to make use of the obtained subunit layouts for in-depth studies of the retinal circuitry. By knowing the subunit locations, targeted stimuli can be manufactured to characterize the spatiotemporal integration properties, nonlinearities and functional synaptic weight of each subunit, and evaluate their similarity across multiple subunits within one ganglion cell receptive field. Interactions between bipolar cells, e.g. via gap junctions, underlying computational functions like motion processing (Kuo et al., 2016; Manookin et al., 2018), could also be investigated more closely by stimulating individual subunits in a specific temporal order. Moreover, since ganglion cell receptive fields tile visual space, one can expect two neighboring ganglion cells to share some of their subunits, corresponding to shared excitatory input (Liu et al., 2017). By studying the same subunit via responses of multiple ganglion cells, ganglion and bipolar cell effects can be disentangled and processes like global and local contrast adaptation (Brown and Masland, 2001; Garvert and Gollisch, 2013; Khani and Gollisch, 2017) be studied in greater detail.

Furthermore, since the number of ganglion cell types generally exceeds the number of bipolar cell types (Masland, 2012), some, if not all, bipolar cell types provide input to multiple ganglion cell types, such that, consequently, the subunit layouts of several ganglion cell types must coincide. If such common subunit layouts could be identified, this would provide a way of studying how amacrine cells shape the signals that bipolar cells send to ganglion cells (Asari and Meister, 2012; Franke et al., 2017). In theory, by identifying all subunit layouts, relating subunit properties to bipolar cell types and studying the transmission strength of each bipolar to ganglion cell type by targeted stimuli, a fairly complete functional connectome of this part of the retina might be in reach, at least as far as connections with sufficiently nonlinear transmission are concerned.

### Potential method extensions

One possible adaptation of the Ricker stripes that might be necessary for some ganglion cell types could be the inclusion of color. For example, the small bistratified ganglion cell in the primate retina is characterized by blue On and yellow Off responses, which are likely conveyed by excitatory input from two types of bipolar cells – a blue-sensitive On bipolar cell and a yellow-sensitive Off bipolar cell (Crook et al. (2009), but see Field et al. (2007)). Both can be expected to form overlapping subunit layouts, but, nevertheless, both could separately be identified by using stimuli with an appropriate chromatic makeup to independently activate different photoreceptor populations (Estévez and Spekreijse, 1982). For example, S-cone-isolating Ricker stripes with an On-type blue center could be applied to reconstruct the blue-sensitive subunits and, conversely, S-cone-silencing stripes with an Off-type center to reconstruct the yellow-sensitive subunits.

Some of the analysis concepts introduced here may also be of interest for studying nonlinear processing beyond the retina. Subunit models have also been applied to primary visual cortex (Vintch et al., 2015; Almasi et al., 2020; Bartsch et al., 2022) and higher motion processing cortical areas (Mineault et al., 2012; Beyeler et al., 2016) as well as to the auditory system (Ahrens et al., 2008; McFarland et al., 2013; Keshishian et al., 2020). Complex cells in the primary visual cortex, for example, display strongly nonlinear response characteristics which can be modeled by subunits that resemble simple cells (Hubel and Wiesel, 1962; Martinez and Alonso, 2003) in a way comparable to the subunits of nonlinear ganglion cells (Carandini et al., 2005). A clearer picture of the organization of these subunits may help understand the functional circuitry of the nonlinear computations in complex cells. For this, the development of suitable visual stimuli may be guided by the idea that the spatial pattern should not necessarily maximize the responses of a subunit but rather sharpen its responses to differentiate it from contributions of other subunits. For a complex cell that prefers edges of a particular orientation, for example, such a stimulus might be a localized, sharp black-white edge whose intensity profile rapidly falls off as one moves away from the black-white transition, or it might comprise an edge with sidebands of opposing contrast on both sides, analogous to the center-surround structure of the Ricker stripes. The desired effect would be to make the response of the complex cell strongly sensitive to the actual positioning of the stimulus relative to the underlying subunit to probe whether the presumed subunit model holds and to potentially identify individual subunits.

More generally, an essential design principle that underlies subunit identification with STR is the application of a stimulus structure that can strongly trigger individual subunits but simultaneously restricts the positions in the probed stimulus space at which responses will occur, so that the effective subunit overlap is reduced. This design principle should be transferrable also to other sensory systems, e.g., by designing auditory stimuli, potentially combined with quasi-tomographic presentation to increase response strength of putative subunits in spectro-temporal space, for identification of functional circuitry underlying nonlinear computations.

## Methods

### Ganglion cell model

In order to study the super-resolved tomographic reconstruction (STR) method in a scenario with known ground truth, we employed an LNLNP model to simulate ganglion cell responses to a flashed presentation of a given stimulus (grayscale image). The model consisted of the following stages: the linear spatial filters that represent the subunits (L), the subunit nonlinearities (N), the weighted linear summation of subunit signals (L), the output nonlinearity (N), and the spike-generating Poisson process (P).

The subunits mark the first linear computation of the stimulus. The simulated visual area was 40 by 40 pixels large and stimulus pixels could attain values from −1 (black) to +1 (white) with 0 corresponding to mean gray. Each subunit was modeled as a 2D Gaussian, whose parameters were two standard-deviation values, an angle of rotation, and the x- and y-position of the center situated in the simulated 40-by-40 pixel space. All subunits were normalized to a volume of unity. In all our simulations, we used positive values for the subunit filters, corresponding to On-type subunits (and, downstream, to simulated On-type ganglion cells.) The generation of subunit layouts is described in the next section. A variation of the Gaussian subunits were cosine-shaped subunits (used where specified), where the value of the subunit depending on the distance from its center reflected a cosine up to its first zero-crossing. In this case, the size of the cosine subunits was chosen such that a Gaussian fitted to them had the required standard deviations and rotation angle.

To calculate the response of the model to a given stimulus, the linear response of each subunit was first computed as a weighted sum of the stimulus pixels, with weights given by the subunit. The linear subunit responses were then passed through the subunit nonlinearities, here typically a half-wave rectification. For one variation, we applied a threshold-quadratic nonlinearity instead, which involved an additional squaring of positive values. Next, the subunit outputs were summed in a weighted manner. In the default setup, all weights were equal, with their sum normalized to unity. For one variation, weights depended on the subunit positions and were determined by the value of a 2D Gaussian centered in the simulated area with a standard deviation of 0.12 times the extent of the area, and, again, were normalized to a sum of unity.

The resulting signal was then transformed into a spike rate by the output nonlinearity, which here amounted to a simple scaling of the signal as rectification was not required due to the already rectified inputs. As reference points, we assumed that background gray (i.e., a signal of zero) would elicit no spikes and that a full-field white flash (i.e., the maximum stimulation) would yield an average response of 30 spikes. Spike rate responses to all other stimuli were determined by linear interpolation between these two reference points. For one variation, we added a universal baseline level of 3 spikes to the spike rate. The resulting spike rate was used as the model output in analyses that were based on noiseless responses. By contrast, when stochastic spike counts were analyzed, the resulting spike rate was converted into an actual spike count using a Poisson process. Here, a single random number was drawn from a Poisson distribution with an expected value given by the spike rate. Thus, the response of the model to a given stimulus was either a spike rate (deterministic) or a random integer spike count (stochastic), depending on the specified model analysis.

The receptive field of a model was empirically determined from responses to individually presented white pixels, with the strength of the receptive field given by the response to the white pixel at the corresponding location. In the default case of rectifying-linear subunit nonlinearities, this noise-free high-resolution measurement corresponds to a weighted sum of the Gaussian subunits.

### Generation of subunit layouts

The biologically inspired subunit layouts used throughout this manuscript were generated using Voronoi diagrams of perturbed hexagonal lattices. To create such a layout, we first constructed a large hexagonal lattice (i.e., the centers of a honeycomb structure) with a nearest-neighbor distance of 1/8^th^ of the simulated area. The points in the lattice were then randomly and independently perturbed by shifting them in x- and y-direction by distances each drawn from a Gaussian distribution with a standard deviation of 21% of the nearest-neighbor distance. Next, we determined the Voronoi cells of all perturbed points using the Euclidean distance and picked those *N* cells (with *N* being the number of desired subunits) whose centers of mass were closest to the center of the lattice. We then fitted 2D Gaussians to these Voronoi cells that we would use as the subunits. The overlap of the subunits was adjusted by multiplying their standard deviations with 1.35 (1.6 as an alternative value used where an increased overlap is specified) and the size of the layout was rendered roughly independent of the number of subunits by scaling with 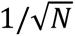 in both spatial directions. In most cases throughout this manuscript, we typically used *N* = 10 subunits. Note that our procedure of fitting Gaussians to Voronoi cells leads to subunits with varying sizes, shapes and rotations.

For the simple subunit layout used in the beginning of this manuscript, four subunits were placed at x-as well as at y-positions of 3/8 and 5/8 of the extent of the simulated area. These Gaussian subunits had standard deviations of 1/10 of the extent of the simulated area in both directions and thus no orientation.

### Stimulus and analysis in simulations

To demonstrate the effect of a suppressive ring around an excitatory spot when probing the receptive field with the spot, we used a stimulus in the shape of a 2D Marr wavelet:

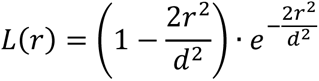

Here, *L*(*r*) is the stimulus intensity (Weber contrast) of a pixel at a distance *r* (in pixels) from the center of the wavelet, and 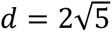 defines the size of the wavelet. The wavelet will take on the maximum allowed intensity of +1 in the center and is normalized to an average intensity of zero. When demonstrating the responses of the model to a spot without a ring, we used the same formula but truncated negative values. These stimuli were centered at every pixel of the simulated area to record responses.

The tomographic stimulus is a stripe with the profile being described by the related 1D Ricker wavelet, hence the name Ricker stripe:

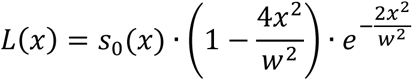

with

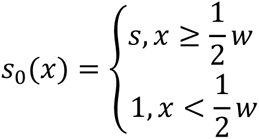

*L*(*x*) is the luminance of a pixel at a distance *x* perpendicular to the center of the Ricker stripe and *w* is the width of the stripe given by the distance of the zero-crossings (i.e., the width of the central white band). The surround factor *s* introduced in our modified definition of the Ricker wavelet only multiplies the negative sidebands of the stripe and affects its integral, which equals zero only if *s* = 1. Again, this wavelet will take on the maximum allowed luminance of +1 in the center. We here applied *w* = 5 pixels *s* = 2.5 if not otherwise specified. If the surround factor *s* would lead to some stimulus pixels attaining values *L* < −1, they are clipped at −1. When demonstrating the principle of the method (Figure 1), the surround factor *s* was set to unity or, in the case without suppressive sidebands, to zero.

Measurements with Ricker stripes were performed at 36 equally distributed angles from 0° (inclusive) to 180° (exclusive). For each angle, 60 equally spaced stripe positions were probed with a shift of 2/3 of a pixel between two positions. Consequently, 2160 combinations of stripe angle and position were tested, with each combination being measured only once, and the resulting responses were composed in a sinogram. If the measurements were performed while using a spiking process, a Gaussian smoothing of the sinogram was undertaken. If not specified otherwise, the applied 2D Gaussian had a standard deviation of *σ*_pos_ = 1.5 stripe positions (corresponding to one pixel of simulated area) in the spatial dimension and *σ*_ang_ = 1.0 angle step sizes (i.e., 5°) in the angle dimension. The resulting sinograms were then reconstructed using FBP with a ramp filter. Note that the resolution of the reconstruction, determined by the distance between stripe positions, is slightly higher than the resolution of the simulated area.

### Calculation of F-scores

To quantify the quality of a reconstruction obtained by FBP, we first identified all local maxima in the reconstruction. We defined a local maximum as a pixel with a value at least as great as any of its eight neighbors. Next, we discarded any local maximum smaller than 30% of the global maximum of the reconstruction. We also discarded any local maximum that lay outside a circle centered in the reconstruction with a diameter of 90% the reconstruction’s extent. We considered all remaining local maxima to be the detected hotspots.

We then identified matches between model subunits and detected hotspots. A hotspot was considered to correspond to a subunit if it lay within the 0.75 σ ellipse of that subunit. The 0.75 σ ellipses never overlapped, and if two hotspots lay within one 0.75 σ ellipse of one subunit, only one hotspot was judged to correspond to the subunit while the other was not considered a match. Consequently, this procedure gave us a number of true positives (hotspots corresponding to subunits), false positives (hotspots not corresponding to a subunit), and false negatives (subunits not detected by any hotspot). As a measure of reconstruction quality, we then calculated the F-score defined as the harmonic mean of precision (true positives over number of hotspots) and sensitivity (true positives over number of subunits):

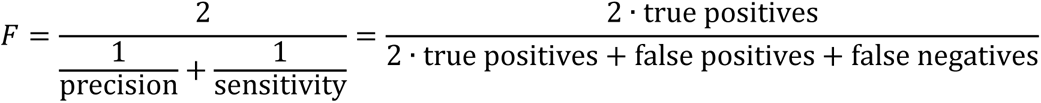

Since the F-score of the reconstruction varies from subunit layout to subunit layout, any numbers given in the text or figures were calculated as averages over 1000 different layout instantiations, resulting in a standard error of the mean below 0.01 in all cases. All F-scores given in text and figures concern models employing a spiking process. Simulations were performed using Python and ran on a desktop computer.

### Experimental procedures

For the experimental test of our method, we used the retina of a 12-year-old adult male marmoset monkey (*Callithrix jacchus*). The retinal tissue was collected directly after the animal was killed for use by other researchers, in conformance with national and institutional guidelines and as approved by the institutional animal care committee of the German Primate Center and by the responsible regional government office (Niedersächsisches Landesamt für Verbraucherschutz und Lebensmittelsicherheit, Permit 33.19-42502-04-20/3458). Following the enucleation procedure, the eyes underwent dissection, during which the cornea, lens, and vitreous humor were extracted to obtain access to the retinal tissue. The retina was then placed in a light-tight chamber that contained Ames’ medium (Sigma-Aldrich, Munich, Germany) supplemented with 6 mM D-glucose and oxygenated with a mixture of 95% O_2_ and 5% CO_2_. To maintain a pH of 7.4, the medium was buffered with 22 mM NaHCO_3_. Following a dark-adaptation period of several hours, during which retinal pieces for other experiments were prepared, a piece of peripheral retina was excised, isolated from the pigment epithelium and placed on a multielectrode array (MultiChannel Systems, Reutlingen, Germany) with 252 electrodes spaced 60 µm apart and sized 10 µm, which had been coated with poly-D-lysine. The entire preparation process was carried out under infrared illumination using a stereomicroscope equipped with night-vision goggles.

While recording from the retina piece, it was continuously supplied with the oxygenated Ames’ medium at a flow rate of 8-9 ml/min. To maintain a stable temperature around 33°C, an inline heater (PH01, MultiChannel Systems, Reutlingen, Germany) and a heating element beneath the array were employed. The recorded multielectrode array signals were amplified and band-pass filtered to a frequency range of 300 Hz to 5 kHz, and saved to disk using the software MC-Rack 4.6.2 (MultiChannel Systems) at a sampling rate of 25 kHz. To identify and sort spikes from the recordings, a modified version of Kilosort (Pachitariu et al., 2016) was used. The original version can be accessed at https://github.com/MouseLand/Kilosort, while the modified version can be found at https://github.com/dimokaramanlis/KiloSortMEA. The output generated by Kilosort was manually reviewed and curated using the software Phy2 (https://github.com/cortex-lab/phy), discarding all units without a distinct cluster of voltage traces or without a clear refractory period.

Custom-made software coded in C++ and OpenGL was utilized to generate visual stimuli. These stimuli were displayed on a gamma-corrected monochromatic white OLED monitor (eMagin) with an 800-by-600 pixel resolution and a refresh rate of 85 Hz. When projected onto the retina through a telecentric lens (Edmund Optics), the pixel size on the retina was 7.5 µm by 7.5 µm. The stimuli described in this manuscript and the gray background illumination between stimuli had a mean irradiance of 5.45 mW/m^2^. We calculated the isomerization rates of the photoreceptors following the formula in Lamb (1995) with literature values for peak sensitivities and collecting areas for marmoset and macaque monkeys (Travis et al., 1988; Schnapf et al., 1990; Tovée et al., 1992; Schneeweis and Schnapf, 1995). This led to 450 isomerizations per photoreceptor per second for S-cones, 2900 for M-cones, and 8600 for rods, indicating a low photopic light regime. The projection of the stimulus screen was focused on the photoreceptors before the start of the experiment, which was confirmed by monitoring through a microscope.

### Basic characterizations of recorded cells

To demonstrate the spatially nonlinear integration of certain ganglion cells, we displayed a reversing-grating stimulus. Square-wave gratings (100% Michelson contrast) with bar widths of 7.5 µm, 15 µm, 30 µm, 60 µm, 120 µm, 240 µm, 480 µm and 6000 µm (full-field) and correspondingly 1, 1, 2, 2, 4, 4, 8, and 1 different spatial phases were presented for 12.5 s each (preceded by 1 s of full-field gray at mean intensity), reversing in contrast every about 0.5 s. The entire sequence was repeated once for a total stimulus duration of about 10 minutes. We calculated PSTHs with a duration of two reversals and a bin size of 10 ms for visualization.

We estimated the receptive fields of cells by computing the spike-triggered average (STA) from responses to a spatiotemporal binary white-noise stimulus on a checkerboard grid (Meister et al., 1994; Chichilnisky, 2001). Each stimulus field had a size of 15 µm by 15 µm and was randomly updated every four frames (i.e., 47 ms) to either black or white with 100% Michelson contrast. The stimulus alternated between regular white noise for 3825 frames (45 s) and a fixed white-noise sequence of 652 frames (∼8 s) for a total time of about one hour. In this study, we only used the non-fixed white noise. The STA was obtained with a temporal window of 42 frames (∼0.5 s). From the spatiotemporal STA, smoothed with a spatial Gaussian filter of 60 µm standard deviation, we then detected the element that had the largest absolute value. We defined the temporal component of the STA as the unsmoothed time course of that pixel and the spatial component as the unsmoothed STA frame of that element. We normalized the temporal component to a Euclidean norm of unity, fitted a 2D Gaussian to the spatial component, and calculated the effective receptive field diameter as the diameter of a circle with the same area as the 1.5 σ ellipse of the Gaussian.

To estimate a ganglion cell’s output nonlinearity, we constructed an LN model, using the spatial and temporal STA components as filters. To enhance signal quality and reduce noise, we truncated the length of the temporal filter to 0.25 seconds. Additionally, only pixels falling within the smallest rectangular window that still contained the 3 σ ellipse of the fitted Gaussian were considered in this computation. Both temporal and spatial filter were normalized to unit Euclidean norm. Applying the temporal and spatial filters to the stimulus yielded a generator signal for each frame of the white-noise stimulus. We grouped the generator signals into ten bins with the same number of data points. The average generator signal of each bin in conjunction with the average of the corresponding spike count gave an estimation of the cell’s contrast-response relationship.

We also computed a 50 ms long autocorrelation function for a cell’s spike train from the non-fixed parts of the white noise stimulus with a resolution of 0.04 ms, smoothed with a Gaussian of 0.4 ms standard deviation, and normalized to a sum of unity.

We manually identified 31 Off and 18 On parasol cells in the dataset, based on their fast biphasic filters, effective receptive field diameters in the expected range of roughly 100 µm to 150 µm (Masri et al., 2019), and tiling of visual space by their receptive fields.

### Stimulus and analysis for subunit identification in experiments

For the application in experiments, the tomographic stimulus was slightly modified in some aspects compared to the analyses of model simulations. While the Ricker stripe profile remained unchanged, we flipped it in polarity so that the center of the stripe was at maximum black (−100% Weber contrast). A surround factor of *s* = 1.5 was applied to the white sidebands, and the width of the Ricker stripes, i.e., the distance of the zero-crossings, was chosen as *w* = 45 µm. Furthermore, in contrast to the simulations, we displayed multiple parallel stripes simultaneously across the entire stimulation area with a center-to-center distance of 375 µm. Stripes were flashed for 153 ms (13 frames) separated by 447 ms (38 frames) of full-field gray. The stripe angles ranged from 0° to 175° in steps of 5°. For each angle, 75 equally spaced shifts of position, ranging from 0 µm to 370 µm with a step size of 5 µm, were applied to the stripes (perpendicular to their orientation), so that the maximum shift was one step less than the stripe distance. Each combination of stripe angle and stripe shift was flashed once in randomized order, before the stimulus was repeated in a new random order. In total, the tomographic stimulus was presented for almost 1.5 hours with each combination presented at least three times.

For each presented combination of stripe angle and position, we determined the average number of spikes elicited between stimulus onset and offset to compose a sinogram. We corrected the sinograms for the positioning of the receptive field by shifting the values in each sinogram row, i.e., for each angle, such that the value at the center corresponded to the stripe position with the smallest distance to the center of the receptive field. Here, we used the position of the 2D Gaussian fitted to the spatial STA from the white-noise analysis as an estimate of the receptive field position. Next, we applied a Gaussian smoothing with a standard deviation of *σ*_pos_ = 1.5 stripe positions (7.5 µm) and *σ*_ang_ = 1 angle step size (5°). The processed sinograms were then used for reconstruction of subunit layouts by applying the same FBP method used for the simulations.

We also analyzed the responses to the offsets of stripes, where we included all spikes during the 153 ms (i.e., the same duration as for the onset) after the offset. Sinogram and FBP analyses were performed analogously. Furthermore, we analyzed the first and second half of the recording separately, by splitting the measurement into the first roughly 45 minutes and second 45 minutes and analyzing both independently.

## Acknowledgments

We thank Fred Rieke for advice on experiments with the primate retina. This work was supported by the European Research Council under the European Union’s Horizon 2020 research and innovation program (grant agreement number 724822), by the Deutsche Forschungsgemeinschaft (German Research Foundation), Projects 515774656, 432680300 (SFB 1456, project B05), and 390729940 (Germany’s Excellence Strategy–EXC 2067/1), and by the Göttingen Graduate Center for Neurosciences, Biophysics, and Molecular Biosciences at the Georg-August-Universität Göttingen.

